# Exotic Tree Plantation Types and Their Distance to Urban Areas in Central Argentina Affect Various Facets of Bird Diversity

**DOI:** 10.64898/2025.12.20.694191

**Authors:** Lucas M. Leveau

## Abstract

Tree plantations are expanding worldwide, negatively affecting native bird communities. However, most research has focused on the impact of tree monocultures on bird taxonomic diversity, while studies on their effects on functional and phylogenetic diversities are limited. This study aimed to compare different bird diversity facets between Eucalypt monocultures and mixed tree plantations, as well as the impact of proximity to urban areas. Bird surveys were conducted during spring using point counts in central Argentina. The mixed tree plantations consisted of *Cedrus deodara, Eucalyptus globulus, Ligustrum lucidum, Pinus taeda,* and *Celtis australis*, with more understorey cover compared to Eucalypt monocultures. Although bird species richness and Simpson diversity were similar between tree plantations, mixed plantations exhibited higher Shannon diversity. Functional and phylogenetic diversities were comparable across plantation types but were significantly influenced by the distance to urban areas. Functional diversity decreased near urban areas, while phylogenetic diversity showed the opposite pattern. The results showed that the increased habitat heterogeneity of mixed plantations likely supported more diverse bird communities, including a greater number of species typical of native woodlands in central Argentina. Additionally, the landscape context significantly influenced the bird communities in tree plantations.

## 1. Introduction

Tree plantations are growing worldwide due to their significance in timber and paper pulp production (Liu et al., 2018; FAO, 2020). As a result, concerns are rising about how tree plantations affect bird diversity compared to natural forests (Schaaf et al., 2024). Several meta-analyses have demonstrated negative impacts of tree plantations on bird diversity and abundance. However, the extent of these impacts was affected by various factors, such as the presence of understory, the origin of the tree species used, and the composition of tree species in plantations (Nájera and Simonetti, 2009; Castaño-Villa et al., 2019; Wang et al., 2022). For example, the presence of understorey in tree plantations can boost habitat diversity, thereby providing more resources to bird species and increasing bird diversity (MacArthur and MacArthur, 1961; Tews et al., 2004; Najera and Simonetti, 2009). Additionally, tree plantations made up of exotic species have shown to have more significant negative effects on bird diversity and abundance compared to plantations of native species (Castaño-Villa et al., 2019; Wang et al., 2022). Furthermore, mixed tree plantations, which consist of multiple tree species, have been shown to support greater bird abundance and diversity than monoculture plantations (Hobson and Bayne, 2000; Kirk and Hobson, 2001; Cavard et al., 2011; but see Jiang et al., 2023).

A deeper understanding of the effects of tree mixed plantations versus tree monocultures can be achieved by analyzing different facets of bird diversity. These aspects include taxonomic diversity, which looks at the number and types of species; functional diversity, which considers the variety and resources used by species; and phylogenetic diversity, which measures how closely related the species are (Díaz and Cabido, 2001; Webb et al., 2002). For example, bird communities can respond differently to types of tree planting depending on whether the focus is on taxonomic, functional, or phylogenetic diversities (Evans et al., 2020; Jayathilake et al., 2021; Schuldt et al., 2022). However, these kinds of analyses are rare worldwide and need more attention.

The impact of tree plantations on bird communities is biased both thematically and geographically, mainly focusing on tree monocultures in tropical regions, while studies on mixed tree plantations are scarce (Castaño-Villa et al., 2019). In particular, in the southern cone of South America, researchers have primarily concentrated on comparing bird communities in Eucalyptus or Pinus plantations and natural habitats (Zurita et al., 2006; Paritsis and Aizen, 2008; Phifer et al., 2017; Jacoboski and Hartz, 2020; Lacoretz et al., 2021).

The Pampa ecoregion in central Argentina was originally made up of grasslands (Mateucci, 2012; Oyarzabal et al., 2018). However, after European colonization, the development of “estancias” or ranches led to the planting of mixed tree plantations for cattle shelter or decoration and monocultures of trees for windbreaks (Mateucci, 2012; Paleo et al., 2016). The effects of different types of tree plantations on bird communities have rarely been studied. Additionally, proximity to urban areas can influence the composition of bird communities in tree plantations. Several studies have identified relationships between bird communities in natural forests and their distance to urban areas (Palomino and Carrascal, 2007; Silvetti et al., 2023). For example, generalist bird species inhabiting urban areas may disperse into nearby forests (Silvetti et al., 2023). However, it is unclear whether proximity to urban areas influences bird communities in mixed and monoculture tree plantations.

Therefore, this study aimed to analyze the taxonomic, functional, and phylogenetic responses of bird communities to different plantation types and their distance to urban areas. I expected higher bird diversity in mixed tree plantations due to their greater habitat diversity. Additionally, proximity to urban areas would influence bird communities in tree plantations because of the spread of urban bird species, such as pigeons and House Sparrows (*Passer domesticus*) (Aronson et al., 2014; La Sorte et al., 2018).

## 2. Methods

### 2.1. Study Area

The study was carried out in tree plantations surrounding two small urban areas of central-east Argentina, Villa Cacique (37°41′00″S, 59°24′00″W, 209 m.a.s.l., 2689 inhabitants) and Barker (37°38′00″S, 59°24′00″W, 248 m.a.s.l., 1241 inhabitants) (Figure 1a). The climate has a mean annual precipitation of 775 mm and a mean annual temperature of 13.3°C (Jaureguy and Bernabe 1987). The study area is located in the Austral Pampas, within the Pampean Phytogeographic Province (Oyarzabal et al. 2018). It presents a pseudosteppe of mesophytes with mountain scrub, represented mainly by *Nassella neesiana, N. trichotoma,* and *Piptochaetium napostense*. Although some native vegetation communities still thrive in the hills (maximum altitude, 450 m.a.s.l.; Pasotti 1958), the landscape is nowadays dominated by crops, pastures, and exotic tree plantations.

**Figure 1.**
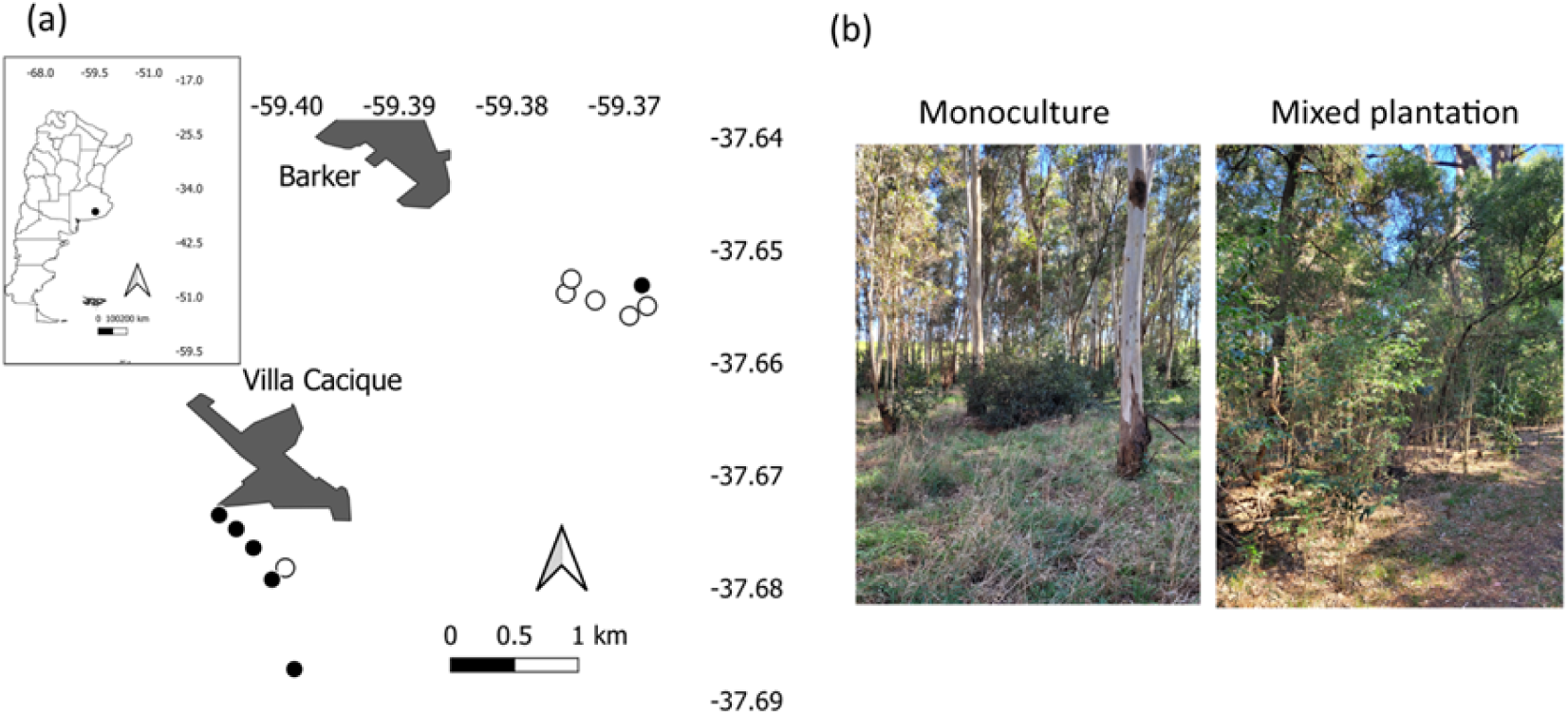
Location of a) study area in Argentina (inset) and survey points in eucalypt monocultures (black dots) and mixed tree plantations (white dots); b) photographs of monoculture and mixed tree plantation. Grey areas indicate urban areas in a).

### 2.2. Tree Plantations

Bird surveys were made in two types of plantations: eucalypt monocultures (hereafter monocultures) and mixed tree plantations (Figure 1b). The plant species were composed mainly of exotic species native to Eurasia, North America, or Australia. Monocultures were dominated by species of the genus Eucalyptus sp. (*E. camaldulensis, E. globulus,* and *E. cinerea*). The undergrowth was dominated by *Dactylis glomerata* and *Laurus nobilis*. The mixed plantations were composed of several tree species, such as *Cedrus deodara, E. globulus, Ligustrum lucidum, Pinus taeda,* and *Celtis australis*, and no single species comprised 80% or more of the total basal area (MacDonald 1995). The undergrowth of the mixed plantations was composed of *C. australis, L. lucidum, L. nobilis, Lonicera fragrantissima, D. glomerata,* and *Dichondra micrantha*, and it had a denser cover than the monocultures (Figure 1b). Moreover, the mixed plantations had fallen dead trees. On the other hand, tree density was higher in monocultures than in mixed plantations (Figure 1b). Eucalypt monocultures and mixed plantations had at least 50 years old.

### 2.3. Bird Surveys

Bird surveys were made using 50-m fixed radius points for 5 minutes, separated by at least 150 m from each other (Bibby et al. 1992). Points were at least 30-50 meters away from the tree plantation boundaries to prevent edge effects (Calviño-Cancela 2013). Birds perching, feeding, or singing were counted, except those flying high over the tree canopy. Surveys were carried out twice during the first four hours after sunrise in the last week of October and mid-December 2024 (austral spring), which coincides with the bird breeding season (de la Peña 2013). Species abundances were averaged between the two visits. Six points were located in each plantation type (Figure 1). The six points of Eucalypt monocultures were spread across three patches: two points in patches of 2.76 and 3.4 hectares, and the other four points in a patch of 11.59 hectares. The six points of mixed tree plantations were located in two patches: one point in a 1.49-hectare patch and the remaining points in a 31.40-hectare patch.

### 2.4. Taxonomic, Functional, and Phylogenetic Diversity

Taxonomic diversity was calculated using Hill’s numbers (Jost 2006). Bird species richness (q = 0) was the number of species seen during the two visits at each point. The Shannon diversity (q = 1) represented the number of common species, whereas the inverse Simpson diversity (q = 2; hereafter Simpson diversity) represented the number of dominant species at each point (Chao et al. 2014). Hill numbers were calculated with the hill_taxa function of the hillR package (Li 2018).

In addition, the accumulated number of species for the three Hill numbers in each plantation type was calculated using rarefaction curves with the online software iNEXT (chao.shinyapps.io/iNEXTOnline). The species accumulation was computed using the species abundances. Based on MacGregor-Fors and Payton (2013), the rarefaction curves were calculated with 999 bootstrap and 84% confidence intervals. Non-overlapping confidence intervals indicated significant differences in species diversity between plantation types.

Functional diversity was calculated using a matrix of species traits related to food resource acquisition (see Leveau 2023). The matrix included the percentage of food types consumed (seeds, fruits, vertebrates, etc.), the percentage use of feeding substrates (ground, canopy, etc.), clutch size (mean number of eggs), body mass (g), and migratory status (Table S1). These data were sourced from Wilman et al. (2014) and de la Peña (2013). A matrix of species functional dissimilarities was constructed using the Gower index with the gowdis function from the FD package in R (Laliberté et al. 2014, R Core Team 2024). The mean pairwise distance of species in each sampling unit was computed using the ses.mpd function from the picante package (Kembel et al. 2010).

This functional diversity index assesses species richness differences between sampling units. I employed the independent swap algorithm (Kembel et al. 2010). Negative values suggest less functional dissimilarity than expected by chance or functional clustering, whereas positive values indicate greater functional dissimilarity than predicted by chance or functional overdispersion. Values close to zero reflect a random functional structure.

The phylogenetic diversity was assessed by downloading a dated phylogeny of bird species from the BirdTree database (https://www.birdtree.org; Jetz et al. 2014). The phylogenetic tree was constructed using TreeAnnotator software (Rambaut and Drummond 2002–2015) and incorporated in R with the read.nexus and as.phylo functions from the ape package (Paradis et al. 2004) (see more details in Leveau 2021). Subsequently, the phylogenetic diversity in each sampling unit was calculated similarly to the functional diversity using the ses.mpd function (Leveau 2021). Negative values indicate greater species relatedness than expected by chance or phylogenetic clustering, while positive values suggest less relatedness than predicted by chance or phylogenetic overdispersion. Values near zero reflect a random phylogenetic structure.

The functional and phylogenetic compositions were assessed using community weighted means (CWMs). These indices were calculated for each trait in every sampling unit with the dbFD function from the FD package in R (Laliberté et al. 2014, R Core Team 2024). The functional traits are those described in Table S1, while the phylogenetic traits represent the species orders (Table 1).

**Table 1.** List of species detected in eucalypt monocultures and mixed tree plantations during spring 2024 in central Argentina. The average number of birds per observation point and the number of occupied points are presented in parentheses (N = 6 for each plantation type). Lines indicate no bird detections. Exotic species are in bold.

| Order | English name | Scientific name | Plantation type |  |
| --- | --- | --- | --- | --- |
|  |  |  | Monoculture | Mixed |
| Falconiformes | Roadside Hawk | *Rupornis magnirostris* | - | 0.08 (1) |
| Falconiformes | Harris's Hawk | *Parabuteo unicinctus* | - | 0.17 (2) |
| Columbiformes | Picazuro Pigeon | *Patagioenas picazuro* | 1.42 (6) | 0.58 (3) |
| Columbiformes | Eared Dove | *Zenaida auriculata* | 0.33 (3) | 0.42 (4) |
| Columbiformes | Picui Ground-Dove | *Columbina picui* | - | 0.75 (3) |
| Apodiformes | Glittering-bellied |  |  |  |
|  | Emerald | *Chlorostilbon lucidus\** | - | 0.33 (2) |
| Apodiformes | White-throated |  |  |  |
|  | Hummingbird | *Leucochloris albicollis* | 0.08 (1) | 0.67 (5) |
| Falconiformes | Chimango Caracara | *Daptrius chimango* | 0.25 (2) | - |
| Psittaciformes | Monk Parakeet | *Myiopsitta monachus* | 1.08 (3) | - |
| Passeriformes | Rufous Hornero | *Furnarius rufus* | 0.92 (4) | 0.75 (4) |
| Passeriformes | Small-billed |  |  |  |
|  | Elaenia | *Elaenia parvirostris\** | 0.08 (1) | 0.42 (4) |
| Passeriformes | White-crested |  |  |  |
|  | Tyrannulet | *Serpophaga subcristata* | 0.75 (4) | 0.58 (5) |
| Passeriformes | Great Kiskadee | *Pitangus sulphuratus* | 0.42 (2) | 0.17 (2) |
| Passeriformes | Tropical Kingbird | *Tyrannus melancholicus\** | 0.25 (2) | 0.25 (2) |
| Passeriformes | Fork-tailed |  |  |  |
|  | Flycatcher | *Tyrannus savana\** | 0.25 (3) | 0.08 (1) |
| Passeriformes | Streaked Flycatcher | *Myiodynastes maculatus\** | 0.08 (1) | 0.08 (1) |
| Passeriformes | Southern House |  |  |  |
|  | Wren | *Troglodytes musculus* | 1.08 (6) | 0.75 (6) |
| Passeriformes | Masked |  |  |  |
|  | Gnatcatcher | *Poliophtila dumicola* | - | 0.08 (1) |
| Passeriformes | Rufous-bellied |  |  |  |
|  | Thrush | *Turdus rufiventris* | 0.33 (3) | 1.42 (6) |
| Passeriformes | Creamy-bellied |  |  |  |
|  | Thrush | *Turdus amaurochalinus* | 0.08 (1) | 0.50 (3) |
| Passeriformes | Tropical Parula | *Setophaga pitayumi* | - | 0.83 (4) |
| Passeriformes | Blue-and-yellow |  |  |  |
|  | Tanager | *Rauenia bonariensis* | - | 0.17 (1) |
| Passeriformes | Double-collared |  |  |  |
|  | Seedeater | *Sporophila caerulescens\** | 0.17 (2) | - |
| Passeriformes | Saffron Finch | *Sicalis flaveola* | 0.42 (4) | 0.42 (3) |
| Passeriformes | Rufous-collared |  |  |  |
|  | Sparrow | *Zonotrichia capensis* | 0.75 (5) | 0.83 (6) |
| Passeriformes | Shiny Cowbird | *Molothrus bonariensis* | 0.08 (1) | - |
| Passeriformes | Screaming Cowbird | *Molothrus rufoaxillaris* | 0.25 (2) | 0.50 (2) |
| Passeriformes | Grayish Baywing | *Agelaioides badius* | 0.42 (4) | 0.25 (2) |
| Passeriformes | Hooded Siskin | *Spinus magellanicus* | 0.17 (2) | 1.58 (5) |
| Passeriformes | **House Sparrow** | ***Passer domesticus*** | 0.58 (3) | 0.25 (1) |
| Total species |  |  | 23 | 26 |

### 2.5. Environmental Variables

The variables explaining bird communities were plantation type (monoculture/mixed) and distance to urban areas. Point counts were at different distances from the urban areas, and proximity to these habitats could influence bird species composition in tree plantations. Therefore, the minimum distance to urban areas (km) was calculated using Google Earth Pro.

### 2.6. Statistical Analyses

The relationship between dependent variables in sampling points and the environmental variables was analyzed using generalized linear models (GLMs). The dependent variables were the taxonomic, functional, and phylogenetic indices. The environmental variables were the plantation types and the distance to urban areas. Due to overdispersion in data, species richness (q = 0) variation was analyzed using the glm.nb function from the MASS package (Ripley et al., 2025). On the other hand, a Gaussian error structure was used for Shannon and Simpson diversities, and CWM values of traits and bird orders. Models were obtained by backward elimination of non-significant variables (P > 0.05) from the full model using the anova function. Final models were compared with null models using a likelihood ratio test (LRT test) (p < 0.05). Multicollinearity among predictor variables was explored using the vif function of the regclass package (Petrie 2020). Multicollinearity was low in our analyses (VIF values < 5). The pseudo-rsquare of final models was obtained using the rsq function of the rsq package (Zhang 2025). Model residuals distribution normality and heteroscedasticity were tested with the DHARMa package (Hartig et al. 2025). Because some points were nested within tree plantation patches, they acted as spatial pseudoreplicates. However, the analysis of spatial autocorrelation of residuals in SAM software (Rangel et al. 2010) showed no significant autocorrelation (Morańs I, P > 0.05). Final model results were plotted with the visreg package (Breheny and Burchett 2017).

The relationship between bird species composition and environmental variables was examined using distance-based Redundancy Analysis (dbRDA). This ordination technique employed species dissimilarity measures between sites to link ordination axes with environmental variables (Legendre and Anderson, 1999). To compare bird assemblages across sites, we used Bray-Curtis (BC) dissimilarity, which accounts for species abundances. dbRDA was performed with the capscale function from the vegan package (Oksanen et al. 2025). Models were developed through backward variable selection and were compared to null models using a likelihood ratio test (LRT) (P < 0.05). In addition, the BC dissimilarity index can be partitioned in components of turnover of individuals and nestedness of individuals between sites (Baselga, 2017). The components of turnover or balanced variation of abundance and nestedness or abundance gradient for all sites were calculated with the beta.multi.abund function of the betapart package in R (Baselga et al., 2025).

## 3. Results

A total of 278 bird detections of 30 species were made (Table 1). Although the accumulated number of species was higher in mixed plantations, no significant differences were found between plantation types (Figure 2). However, the accumulated richness of common species was significantly higher in mixed plantations than Eucalypt monocultures (Figure 2). The number of dominant species was higher in mixed plantations, although differences between plantation types were not significant (Figure 2).

**Figure 2.**
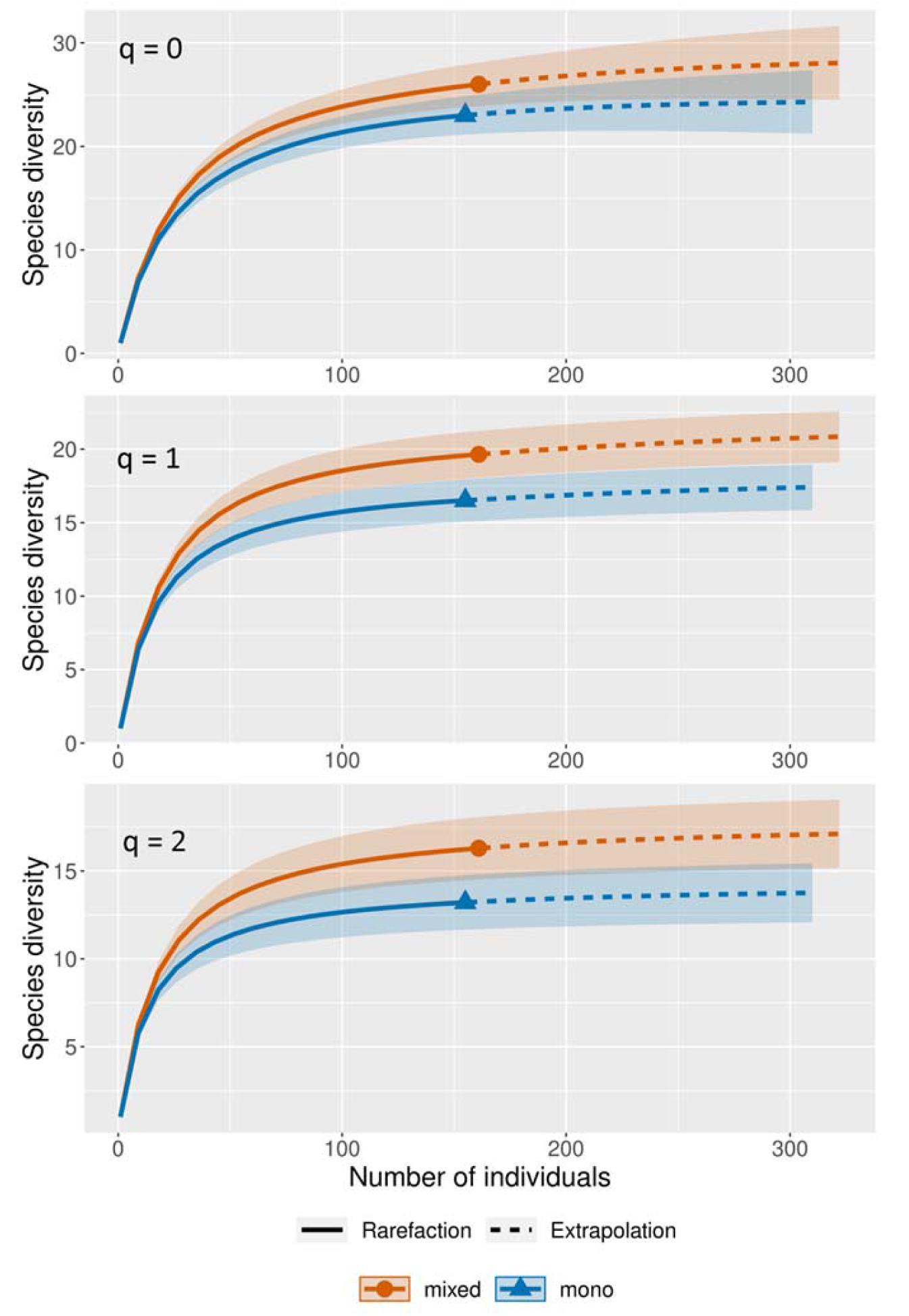
Rarefaction curves of Hill numbers (species richness, q = 0; Shannon diversity, q = 1; and Simpson diversity, q = 2) with sample coverage for mixed tree plantations (mixed) and eucalypt monocultures (mono) during spring 2024, central Argentina. The shaded bands represent the 84% confidence intervals.

At the point scale (N = 12), the mean values of species richness, Shannon diversity, and Simpson diversity were 12.08 (Standard deviation = 4.10), 10.49 (SD = 3.37), and 9.19 (SD = 2.76), respectively. No significant differences in bird richness, Shannon diversity or Simpson diversity were found between plantation types or the distance to urban areas (Table 2).

**Table 2.** Final generalized linear models for bird species richness, Shannon diversity, and Simpson diversity in tree plantations of central Argentina. SES: standardized effect sizes.

| Response variable | Parameter | Estimate | Std. Error | z value | P |
| --- | --- | --- | --- | --- | --- |
| Bird richness ($q = 0$) | Intercept | 2.492 | 0.093 | 26.860 | <0.001 |
| Shannon diversity ($q = 1$) | Intercept | 10.494 | 0.972 | 10.800 | <0.001 |
| Simpson diversity ($q = 2$) | Intercept | 9.186 | 0.796 | 11.540 | <0.001 |
| Functional diversity |  |  |  |  |  |
| (SES) | Intercept | -1.127 | 0.519 | -2.171 | 0.055 |
|  | Urban distance | 1.130 | 0.433 | 2.612 | 0.026 |
| Phylogenetic diversity |  |  |  |  |  |
| (SES) | Intercept | 0.816 | 0.300 | 2.718 | 0.022 |
|  | Urban distance | -0.987 | 0.250 | -3.947 | 0.003 |

The partitioning of total species dissimilarity (BC = 0.83) showed that balanced variation of abundance was the predominant pattern (0.76 vs 0.08 of abundance gradient). Therefore, the difference in species composition across sampling points was mainly caused by a turnover of individuals from different species. The dbRDA showed that two axes account for 34% of the total variation in species composition across sampling points. Plantation type and urban distance explained significantly the variation in species composition (LRT = 2.27, P = 0.004; Figure 3). *Spinus magellanica, Turdus rufiventris,* and *Setophaga pitiayumi* were more abundant in mixed plantations (Figure 3, Table 1). *Patagioenas picazuro, Myiopsitta monachus,* and *Passer domesticus* were more abundant in monoculture plantations and near urban areas (Figure 3, Table 1).

**Figure 3.**
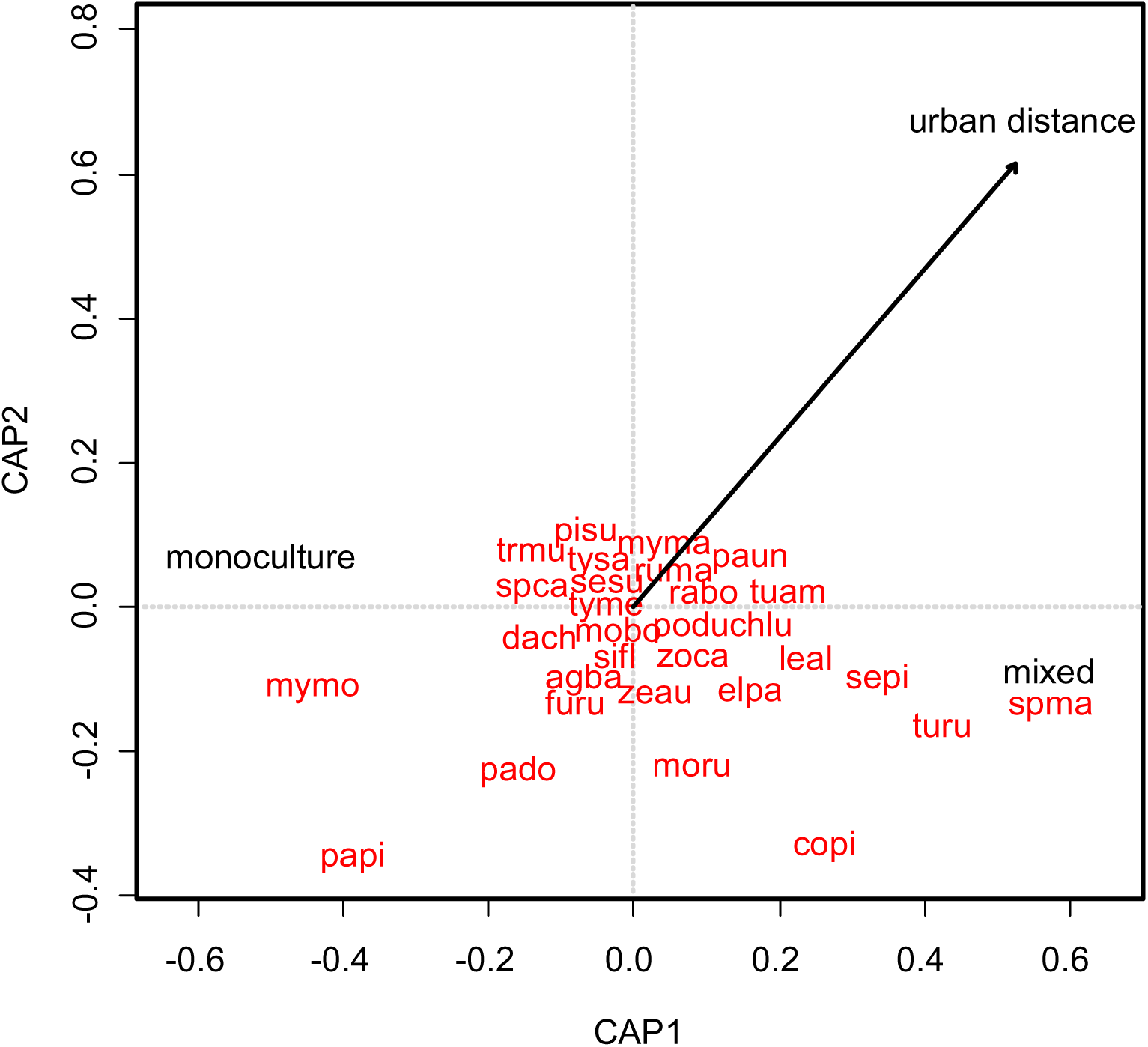
Distance-based redundancy analysis showing the relationship between the environmental variables and bird species composition in tree plantations of central Argentina. monoculture: eucalypt monocultures; mixed: mixed tree plantations; urban distance: minimum distance to urban areas (km). Species names in alphabetical order: agba, *Agelaioides badius*; chlu, *Chlorostilbon lucidus*; copi, *Columbina picui*; dach, *Daptrius chimango*; elpa, *Elaenia parvirostris*; furu, *Furnarius rufus*; leal, *Leucochloris albicollis*; mobo, *Molothrus bonariensis*; moru, *Molothrus rufoaxillaris*; myma, *Myiodynastes maculatus*; mymo, *Myiopsitta monachus*; paun, *Parabuteo unicinctus*; pado, *Passer domesticus*; papi, *Patagioenas picazuro*; pisu, *Pitangus sulphuratus*; podu, *Polioptila dumicola*; rabo, *Rauenia bonariensis*; ruma, *Rupornis magnirostris*; sesu, *Serpophaga subcristata*; sepi, *Setophaga pitiayumi*; sifl, *Sicalis flaveola*; spma, *Spinus magellanicus*; spca, *Sporophila caerulescens*; trmu, *Troglodytes muscuclus*; tuam, *Turdus amaurochalinus*; turu, *Turdus rufiventris*; tyme, *Tyrannus melancholicus*; tysa, *Tyrannus savana*; zeau, *Zenaida auriculata*; zoca, *Zonotrichia capensis*.

The SES of functional and phylogenetic diversities in tree plantations were significantly related to the minimum distance to urban areas (LRT = 7.91, P = 0.009, r^2^ = 0.41; LRT = 6.03, P < 0.001, r^2^ = 0.61, respectively) (Table 2). While functional SES values decreased with urban proximity (Figure 4), the phylogenetic SES values increased. Therefore, plantations near urban areas had functional clustered assemblages belonging to different phylogenetic clades.

**Figure 4.**
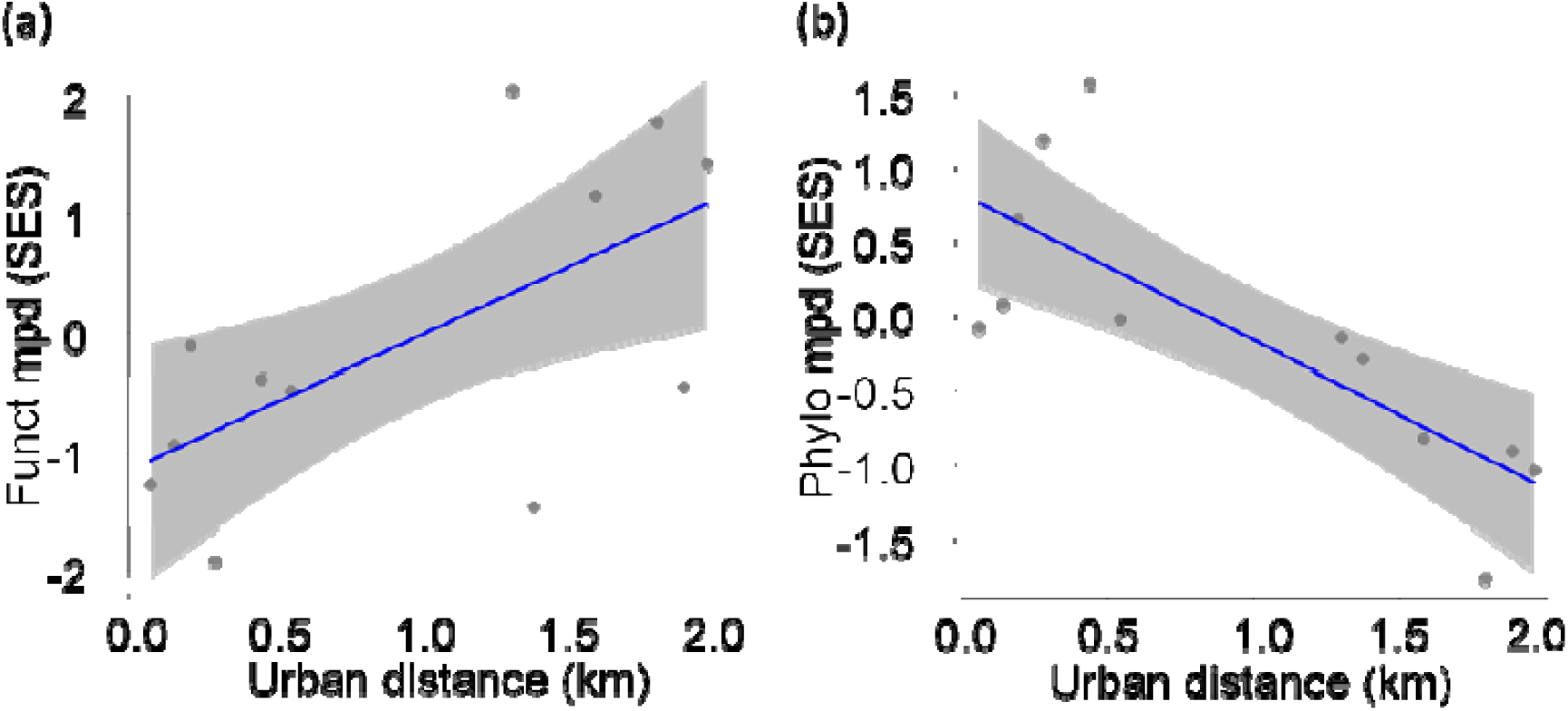
Relationship between the minimum distance to urban areas and the standardized effect sizes (SES) of a) functional diversity (Funct mpd) and b) phylogenetic diversity (Phylo mpd) in point counts (N = 12) of tree plantations in central-east Argentina during spring 2024. Blue lines are fitted models and grey bands are 95% confidence intervals.

Functional and phylogenetic composition were related to plantation types and distance from urban areas (Tables S1 and S2). The abundance of nectarivorous birds was higher in mixed plantations (Figure 5a), while species that feed on the understory and have larger body mass were more abundant in monocultures (Figures 5d, and 5e). The number of granivorous birds increased with proximity to urban areas (Figure 5b), while the abundance of understory species tended to grew (P = 0.076) as one moved further from urban areas (Figure 5c).

**Figure 5.**
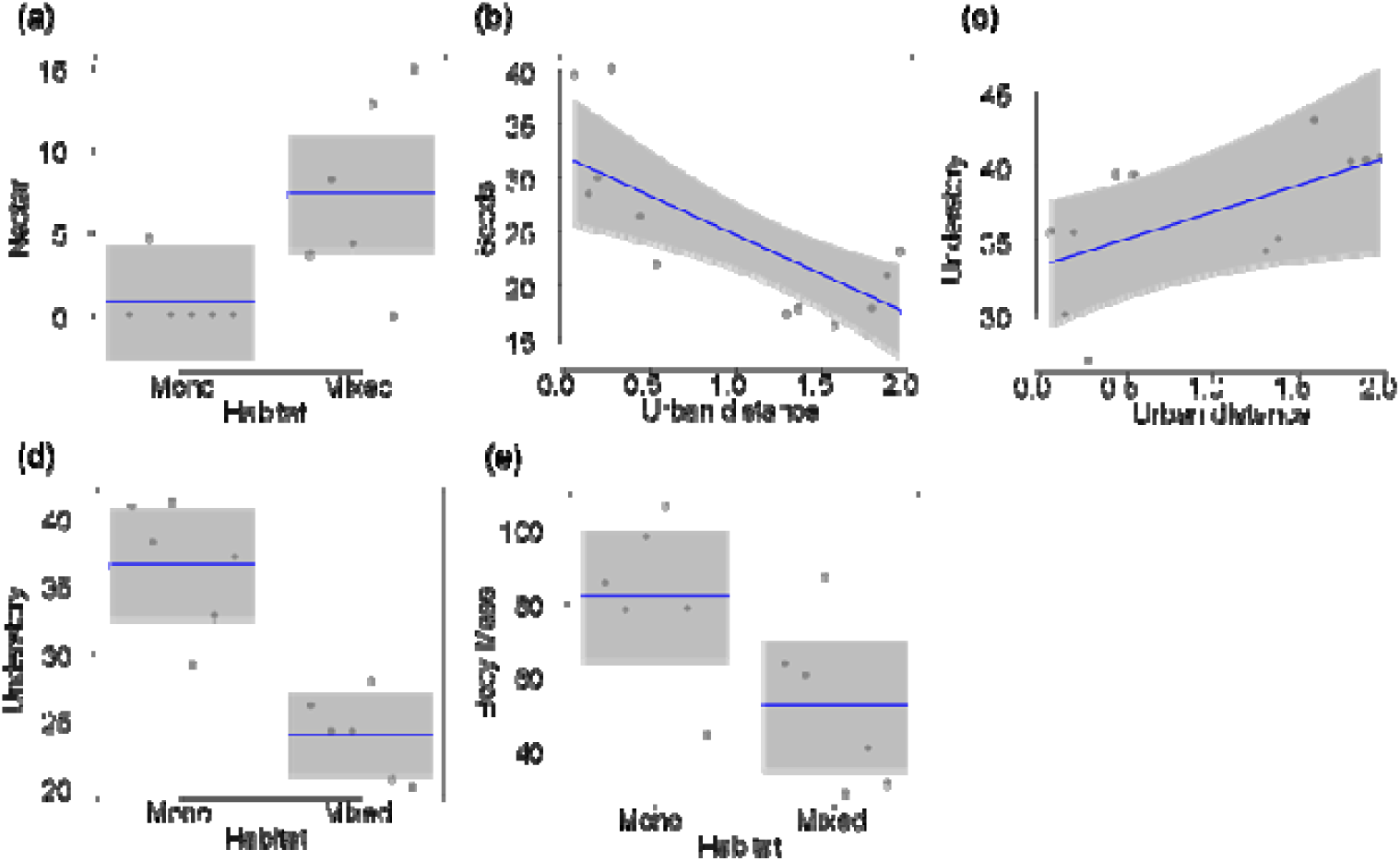
Relationship between the community weighted means of functional traits and environmental variables of tree plantations in central-east Argentina. Mono: eucalypt monocultures; Mixed: mixed tree plantations. Blue lines are fitted models and means, and grey bands are 95% confidence intervals.

Monocultures had a higher abundance of species belonging to Apodiformes (Figure 6a). Proximity to urban areas was positively related to the abundance of Columbiformes (Figure 6b). In contrast, the abundance of Passeriformes increased with the distance to urban areas (Figure 6c).

**Figure 6.**
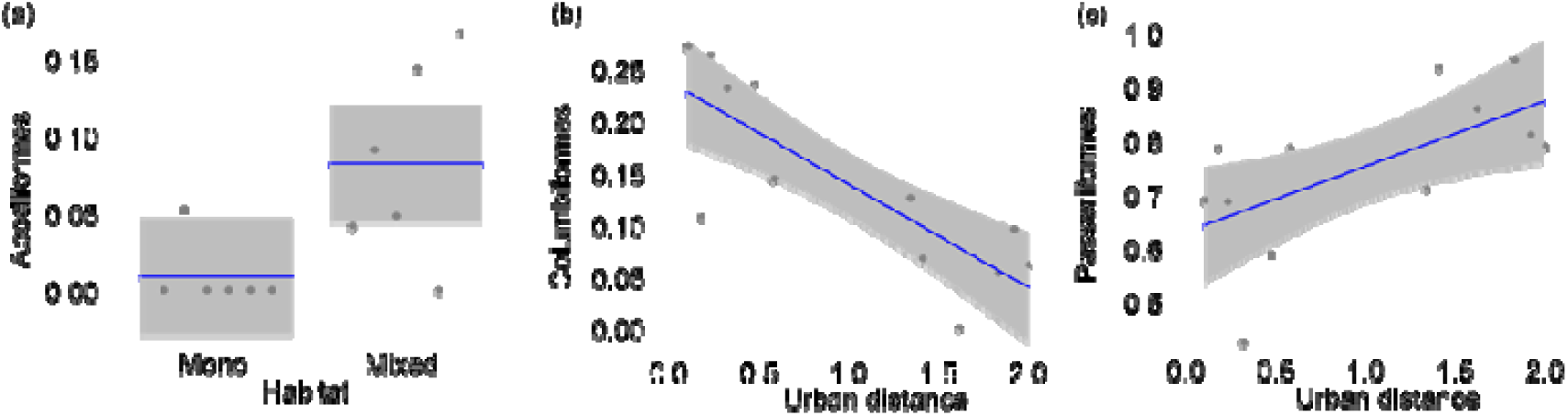
Relationship between the community weighted means of bird orders and environmental variables of tree plantations in central-east Argentina. Mono: eucalypt monocultures; Mixed: mixed tree plantations. Blue lines are fitted models and means, and grey bands are 95% confidence intervals.

## 4. Discussion

The results showed significant differences in bird communities between the different types of tree plantations. Mixed tree plantations supported greater bird diversity than monocultures. Additionally, the functional and phylogenetic diversity of sites was linked to the distance from urban areas, indicating that the landscape context of tree plantations influences community assembly processes.

### 4.1. Patterns of Bird Taxonomic Diversity and Composition

Although there were no differences in species richness or diversity at the sampling point level between plantation types, the total richness of common species was higher in mixed plantations than in eucalypt plantations. This pattern supports our habitat diversity hypothesis, suggesting that mixed tree plantations may provide more resources and microhabitats for a greater number of bird species. On one hand, different tree species offer various food resources, nesting sites, and shelter for bird species compared to eucalypt plantations. For example, some insectivorous birds such as *Polioptila dumicola* and *Setophaga pitiayumi* have been observed foraging in coniferous trees like *Cedrus deodara* and *Pinus* sp. (L.M. Leveau, pers. obs., Leveau, 2023). On the other hand, the presence of understorey and dead trees in mixed plantations can provide additional nesting and food resources (Fuller et al. 2012, Wesołowski et al. 2018). For instance, frugivorous birds such as *Turdus rufiventris* and *Turdus amaurochalinus* can feed on the fruits of understorey trees like *Celtis australis*.

Although these plantation types were established in the Pampa grasslands, they could provide habitat for bird species typical of native coastal woodlands of central Argentina. Mixed tree plantations supported greater numbers of *Turdus rufiventris, Turdus amaurochalinus, Setophaga pitiayumi, Elaenia parvirostris, Polioptila dumicola,* and *Rauenia bonariensis*, species considered woodland-dependent (Cueto and Lopez de Casenave, 2000; Horlent et al., 2003; Lacoretz et al., 2021; Godoy et al., 2024). Several of these species, such as *Turdus amaurochalinus, Setophaga pitiayumi,* and *Elaenia parvirostris*, have not been recorded in the study area for at least 25 years (L.M. Leveau, obs. pers.), indicating a process of range expansion and increasing abundance in central Argentina (Galiano et al. 2024; Leveau, 2024).

Bird species composition in tree plantations was related to the distance from urban areas. Tree plantations near urban areas had higher abundances of *Patagioenas picazuro, Myiopsitta monacha,* and *Passer domesticus*. These species are common and abundant in urban areas of central Argentina because of lawned areas that provide food resources (Leveau 2019, Galiano et al. 2024). For *Passer domesticus*, buildings would offer nesting sites (Lowther and Cink, 2020). Therefore, proximity to urban areas may facilitate the spread of these bird species into tree plantations.

### 4.2. Patterns of Functional and Phylogenetic Diversity

Proximity to urban areas and plantation type were significantly related to functional and phylogenetic diversities. Tree plantations near urban areas showed lower functional diversity than expected by chance. Bird species in these areas had more similar functional traits compared to those far from urban regions. This pattern aligns with studies conducted in cities, where species are filtered based on their functional traits (Weideman et al. 2020; Leveau 2021; Melo et al. 2022; Moreno-Contreras et al., 2024; He et al. 2025). Urban areas are primarily occupied by granivorous bird species that feed on the ground, while non-urban areas are inhabited by more species with insectivorous diets that feed in the canopy or understorey (La Sorte et al., 2018; Ordóñez-Delgado et al., 2022; Melo et al., 2022; Leveau, 2022). Therefore, proximity to urban areas promoted the dominance of granivorous bird species in tree plantations. In contrast, tree plantations located farther from urban areas had a higher abundance of bird species feeding on the understorey.

Conversely, the phylogenetic diversity of bird communities in tree plantations increased closer to urban areas. Therefore, bird species were less phylogenetically related than those in bird communities farther from urban areas. This pattern matches findings from central Argentina, where urban bird communities tended to be randomly related and rural bird communities were phylogenetically clustered (Leveau, 2021). Urban bird communities tend to favor the presence and abundance of Columbiformes (La Sorte et al. 2018; Leveau, 2022; Moreno-Contreras et al., 2024), thus adding distant clades to other common groups such as Passeriformes and increasing the phylogenetic diversity of communities. Therefore, tree plantations farther from urban areas were dominated by Passeriformes species, while plantations near urban areas also included Columbiformes species. The proximity of urban areas probably promoted the dispersal of Columbiformes into tree plantations, boosting their phylogenetic diversity.

The combined findings on the functional and phylogenetic traits of bird communities indicate their assembly process in relation to urban proximity. Bird communities near urban areas consist of species with similar functional traits but that are phylogenetically distant. Therefore, the assembly of these bird communities appears to be a convergent selection of distant species with similar traits that help them persist in urban areas. Conversely, bird communities in tree plantations far from urban areas had species with different functional traits but that are phylogenetically related (see also Leveau 2021).

The functional and phylogenetic composition was linked to the plantation type. Bird species of Apodiformes that feed on nectar were more abundant in mixed tree plantations. These species, such as *Chlorostilbon aureoventris* and *Leucochloris albicollis*, could access more floral resources in the more diverse understorey of the mixed tree plantations. Additionally, mixed plantations could provide adequate microhabitats and materials for nesting, since these species need mosses and lichens to build their nests (Povedano y Maugeri, 2020). Counterintuitively, eucalypt monocultures had a higher abundance of birds that feed on the understorey. *Troglodytes musculus*, which was more abundant in monocultures, probably benefited from the availability and abundance of food resources. On the other hand, bird species in monocultures tended to have larger body sizes compared to those in mixed tree plantations. This pattern was likely due to the greater abundance of pigeons and parrots, such as *Patagioenas picazuro* and *Myiopsitta monachus*, in monocultures.

### 4.3. Study Limitations

The findings of this study should be interpreted with caution due to several issues. First, the sample size was small, and future efforts should include more tree plantations. Second, several environmental variables that could influence the patterns of bird communities were not measured in this study. For example, soil type, tree age, foliage density, bare ground cover, and nesting and food resources are important factors shaping bird communities in forests (Fuller et al. 2012). They should be considered in future studies. Lastly, bird communities in tree plantations may experience seasonal and yearly changes (Wesołowski et al. 2018) that could alter the results of this study.

## 5. Conclusions

The results showed that mixed tree plantations had more bird diversity than eucalypt plantations, and bird composition changed significantly according to plantation type and distance to urban areas. Therefore, dispersion and habitat selection probably shaped bird communities. The species turnover between plantation types was perhaps related to the different habitat requirements of bird species. Additionally, bird community assembly was mainly linked to proximity to urban areas. Tree plantations near cities hosted bird species that were functionally similar but phylogenetically diverse. However, more research is needed to disentangle the roles of plantation age, soil type, and food and nesting resources influencing bird communities.

Despite the limitations of this study, some recommendations for tree plantation management are suggested. First, to promote the presence of understorey vegetation and dead trees, which can provide resources for bird species. Second, to encourage the use of mixed tree plantations for commercial purposes, which can also enhance bird diversity.

## Notes

### Competing Interest Statement

The authors have declared no competing interest.

