## Supplementary material for "Exotic Tree Plantation Types and Their Distance to Urban Areas in Central Argentina Affect Various Facets of Bird Diversity": Table S1, Table S2

Table S1. Functional traits of species. For more information see Wilman et al. (2014). Species names are in alphabetical order.

|  |  | Diet | | | | | | | | | Substrate | | | |  |  |  |
| --- | --- | --- | --- | --- | --- | --- | --- | --- | --- | --- | --- | --- | --- | --- | --- | --- | --- |
| Species | Clutch | Inv | Vend | Vect | Vfish | Scav | Fruit | Nect | Seed | PlantO | Ground | Understory | Midhigh | Canopy | BodyMass | Migrant | Resident |
| *Agelaioides badius* | 6.5 | 60 | 0 | 0 | 0 | 0 | 0 | 0 | 40 | 0 | 50 | 30 | 20 | 0 | 45.25 | 0 | 1 |
| *Chlorostilbon lucidus* | 2 | 10 | 0 | 0 | 0 | 0 | 0 | 90 | 0 | 0 | 0 | 0 | 90 | 10 | 3.5 | 1 | 0 |
| *Columbina picui* | 2 | 0 | 0 | 0 | 0 | 0 | 0 | 0 | 100 | 0 | 100 | 0 | 0 | 0 | 47 | 0 | 1 |
| *Daptrius chimango* | 2.5 | 20 | 20 | 10 | 0 | 50 | 0 | 0 | 0 | 0 | 100 | 0 | 0 | 0 | 296 | 0 | 1 |
| *Elaenia parvirostris* | 2.5 | 70 | 0 | 0 | 0 | 0 | 30 | 0 | 0 | 0 | 0 | 33 | 33 | 33 | 13.8 | 1 | 0 |
| *Furnarius rufus* | 3 | 80 | 0 | 0 | 0 | 0 | 0 | 0 | 20 | 0 | 70 | 30 | 0 | 0 | 46.42 | 0 | 1 |
| *Leucochloris albicollis* | 2 | 10 | 0 | 0 | 0 | 0 | 0 | 90 | 0 | 0 | 0 | 30 | 60 | 10 | 6.25 | 0 | 1 |
| *Molothrus bonariensis* | 10 | 70 | 0 | 0 | 0 | 0 | 0 | 0 | 30 | 0 | 100 | 0 | 0 | 0 | 41.49 | 0 | 1 |
| *Molothrus rufoaxillaris* | 9.5 | 30 | 0 | 0 | 0 | 0 | 10 | 0 | 60 | 0 | 90 | 10 | 0 | 0 | 47.5 | 0 | 1 |
| *Myiodinastes maculatus* | 3 | 40 | 0 | 30 | 0 | 0 | 30 | 0 | 0 | 0 | 0 | 0 | 100 | 0 | 43,2 | 1 | 0 |
| *Myiopsitta monachus* | 6 | 10 | 0 | 0 | 0 | 0 | 30 | 0 | 30 | 30 | 20 | 20 | 40 | 20 | 120 | 0 | 1 |
| *Passer domesticus* | 3.5 | 10 | 0 | 0 | 0 | 0 | 0 | 0 | 60 | 30 | 50 | 50 | 0 | 0 | 26.51 | 0 | 1 |
| *Patagioenas picazuro* | 1 | 10 | 0 | 0 | 0 | 0 | 30 | 0 | 30 | 30 | 30 | 30 | 30 | 10 | 279 | 0 | 1 |
| *Parabuteo unicinctus* | 2 | 0 | 90 | 10 | 0 | 0 | 0 | 0 | 0 | 0 | 0 | 100 | 0 | 0 | 850.28 | 0 | 1 |
| *Pitangus sulphuratus* | 3.5 | 40 | 10 | 10 | 10 | 0 | 30 | 0 | 0 | 0 | 50 | 40 | 10 | 0 | 62.85 | 0 | 1 |
| *Polioptila dumicola* | 3 | 100 | 0 | 0 | 0 | 0 | 0 | 0 | 0 | 0 | 0 | 0 | 20 | 80 | 7 | 0 | 1 |
| *Rauenia bonariensis* | 3 | 0 | 0 | 0 | 0 | 0 | 80 | 0 | 10 | 10 | 0 | 30 | 30 | 40 | 36 | 0 | 1 |
| *Rupornis magnirostris* | 3 | 40 | 30 | 30 | 0 | 0 | 0 | 0 | 0 | 0 | 33 | 33 | 33 | 0 | 269 | 0 | 1 |
| *Setophaga pitiayumi* | 2 | 70 | 0 | 0 | 0 | 0 | 20 | 0 | 0 | 10 | 0 | 0 | 0 | 100 | 6.82 | 0 | 1 |
| *Serpophaga subcristata* | 2 | 100 | 0 | 0 | 0 | 0 | 0 | 0 | 0 | 0 | 0 | 33 | 33 | 33 | 6.6 | 0 | 1 |
| *Sicalis flaveola* | 3 | 0 | 0 | 0 | 0 | 0 | 0 | 0 | 100 | 0 | 40 | 60 | 0 | 0 | 16.89 | 0 | 1 |
| *Sporophila caeruslescens* | 3 | 20 | 0 | 0 | 0 | 0 | 0 | 0 | 80 | 0 | 50 | 50 | 0 | 0 | 9.73 | 1 | 0 |
| *Spinus magellanicus* | 3.5 | 10 | 0 | 0 | 0 | 0 | 0 | 0 | 30 | 60 | 33 | 33 | 33 | 0 | 13.6 | 0 | 1 |
| *Troglodytes musculus* | 6 | 80 | 0 | 0 | 0 | 0 | 0 | 0 | 0 | 20 | 0 | 100 | 0 | 0 | 10.85 | 0 | 1 |
| *Turdus amaurochalinus* |  | 40 | 0 | 0 | 0 | 0 | 60 | 0 | 0 | 0 | 20 | 20 | 30 | 30 | 57,9 | 0 | 1 |
| *Turdus rufiventris* | 4 | 50 | 0 | 0 | 0 | 0 | 50 | 0 | 0 | 0 | 100 | 0 | 0 | 0 | 69.44 | 0 | 1 |
| *Tyrannus melancholicus* | 2.5 | 100 | 0 | 0 | 0 | 0 | 0 | 0 | 0 | 0 | 0 | 0 | 50 | 50 | 37.4 | 1 | 0 |
| *Tyrannus savana* | 2.5 | 70 | 0 | 0 | 0 | 0 | 30 | 0 | 0 | 0 | 50 | 50 | 0 | 0 | 31.9 | 1 | 0 |
| *Zenaida auriculata* | 2 | 0 | 0 | 0 | 0 | 0 | 0 | 0 | 100 | 0 | 20 | 20 | 40 | 20 | 110.2 | 0 | 1 |
| *Zonotrichia capensis* | 2.5 | 30 | 0 | 0 | 0 | 0 | 0 | 0 | 50 | 20 | 100 | 0 | 0 | 0 | 20.31 | 0 | 1 |

Table S2. Final generalized linear models between community weighted means of functional traits and environmental variables of tree plantations in central Argentina. LRT: final models compared to null models using a likelihood ratio test (LRT).

| Trait | Parameter | Estimate | Std. Error | t value | P | LRT | P |
| --- | --- | --- | --- | --- | --- | --- | --- |
| Clutch size | Intercept | 3.406 | 0.118 | 28.940 | <0.001 | 0.404 | 0.092 |
| Invertebrates | Intercept | 41.923 | 3.423 | 12.250 | <0.001 | 185.620 | 0.243 |
| Vertebrates | Intercept | 2.880 | 0.773 | 3.725 | 0.003 | 30.379 | 0.060 |
| Fruits | Intercept | 13.299 | 1.011 | 13.15 | <0.001 | 10.766 | 0.352 |
| Nectar | Intercept | 0.790 | 1.754 | 0.450 | 0.662 | 128.960 | 0.008 |
|  | Habitat: mixed | 6.556 | 2.481 | 2.643 | 0.025 |  |  |
| Seeds | Intercept | 31.957 | 3.112 | 10.269 | <0.001 | 11.285 | <0.001 |
|  | Distance | -7.286 | 2.197 | -3.316 | 0.008 |  |  |
| Plant | Intercept | 12.707 | 1.338 | 9.495 | <0.001 | 27.928 | 0.247 |
| Ground | Intercept | 40.900 | 2.700 | 15.150 | <0.001 | 51.773 | 0.451 |
| Understory | Intercept | 33.251 | 2.233 | 14.890 | <0.001 | 13.138 | 0.001 |
|  | Habitat: mixed | -12.693 | 2.792 | -4.545 | 0.001 |  |  |
|  | Distance | 3.642 | 1.814 | 2.008 | 0.076 |  |  |
| Midhigh | Intercept | 17.238 | 1.741 | 9.901 | <0.001 | 9.913 | 0.892 |
| Canopy | Intercept | 11.309 | 1.645 | 6.876 | <0.001 | 118.380 | 0.107 |
| Body mass | Intercept | 81.946 | 9.043 | 9.062 | <0.001 | 2715.000 | 0.019 |
|  | Habitat: mixed | -30.083 | 12.789 | -2.352 | 0.041 |  |  |
| Migrant | Intercept | 0.082 | 0.024 | 3.352 | 0.006 | 0.012 | 0.177 |

Table S3. Final generalized linear models between community weighted means of avian orders and environmental variables of tree plantations in central Argentina.

| Order | Parameter | Estimate | Std. Error | t value | P | LRT | P |
| --- | --- | --- | --- | --- | --- | --- | --- |
| Apodiformes | Intercept | 0.009 | 0.019 | 0.450 | 0.662 | 0.016 | 0.008 |
|  | Habitat: mixed | 0.073 | 0.028 | 2.643 | 0.025 |  |  |
| Columbiformes | Intercept | 0.233 | 0.027 | 8.667 | <0.001 | 0.061 | <0.001 |
|  | Distance | -0.099 | 0.022 | -4.427 | 0.001 |  |  |
| Falconiformes | Intercept | 0.021 | 0.009 | 2.303 | 0.042 | <0.001 | 0.761 |
| Passeriformes | Intercept | 0.633 | 0.058 | 10.977 | <0.001 | 0.092 | 0.011 |
|  | Distance | 0.122 | 0.048 | 2.531 | 0.030 |  |  |
| Psittaciformes | Intercept | 0.047 | 0.031 | 1.504 | 0.161 | 0.026 | 0.108 |
